# Integrating Host Response and Unbiased Microbe Detection for Lower Respiratory Tract Infection Diagnosis in Critically Ill Adults

**DOI:** 10.1101/375360

**Authors:** Charles Langelier, Katrina L Kalantar, Farzad Moazed, Michael R. Wilson, Emily D. Crawford, Thomas Deiss, Annika Belzer, Samaneh Bolourchi, Saharai Caldera, Monica Fung, Alejandra Jauregui, Katherine Malcolm, Amy Lyden, Lillian Khan, Kathryn Vessel, Jenai Quan, Matt Zinter, Charles Y. Chiu, Eric D. Chow, Jenny Wilson, Steve Miller, Michael A. Matthay, Katherine S. Pollard, Stephanie Christenson, Carolyn S. Calfee, Joseph L. DeRisi

## Abstract

Lower respiratory tract infections (LRTI) lead to more deaths each year than any other infectious disease category. Despite this, etiologic LRTI pathogens are infrequently identified due to limitations of existing microbiologic tests. In critically ill patients, non-infectious inflammatory syndromes resembling LRTI further complicate diagnosis. To address the need for improved LRTI diagnostics, we performed metagenomic next-generation sequencing (mNGS) on tracheal aspirates from 92 adults with acute respiratory failure and simultaneously assessed pathogens, the airway microbiome and the host transcriptome. To differentiate pathogens from respiratory commensals, we developed rules-based and logistic regression models (RBM, LRM) in a derivation cohort of 20 patients with LRTI or non-infectious acute respiratory illnesses. When tested in an independent validation cohort of 24 patients, both models achieved accuracies of 95.5%. We next developed pathogen, microbiome diversity, and host gene expression metrics to identify LRTI-positive patients and differentiate them from critically ill controls with non-infectious acute respiratory illnesses. When tested in the validation cohort, the pathogen metric performed with an AUC of 0.96 (95% CI = 0.86 - 1.00), the diversity metric with an AUC of 0.80 (95% CI = 0.63 – 0.98), and the host transcriptional classifier with an AUC of 0.88 (95% CI = 0.75 – 1.00). Combining these achieved a negative predictive value of 100%. This study suggests that a single streamlined protocol offering an integrated genomic portrait of pathogen, microbiome and host transcriptome may hold promise as a novel tool for LRTI diagnosis.

**SIGNIFICANCE STATEMENT:** Lower respiratory tract infections (LRTI) are the leading cause of infectious disease-related death worldwide yet remain challenging to diagnose because of limitations in existing microbiologic tests. In critically ill patients, non-infectious respiratory syndromes that resemble LRTI further complicate diagnosis and confound targeted treatment. To address this, we developed a novel metagenomic sequencing-based approach that simultaneously interrogates three core elements of acute airway infections: the pathogen, airway microbiome and host response. We studied this approach in a prospective cohort of critically ill patients with acute respiratory failure and found that combining pathogen, microbiome and host gene expression metrics achieved accurate LRTI diagnosis and identified etiologic pathogens in patients with clinically identified infections but otherwise negative testing.

## INTRODUCTION

Lower respiratory tract infections (LRTI) are a leading cause of mortality worldwide(1–3). Early and accurate determination of acute respiratory disease etiology is crucial for implementing effective pathogen-targeted therapies but is often not possible due to the limitations of current microbiologic tests in terms of sensitivity, speed, and spectrum of available assay targets(4). For instance, even with the best available clinical diagnostics, a contributory pathogen can be detected in only 38% of adults with community acquired pneumonia, due to the low sensitivity and time requirements of culture, and the limited number of microbes detectable by serologic and polymerase chain reaction (PCR) assays(4, 5).

In the absence of a definitive microbiologic diagnosis, clinicians may presume symptoms are due to a non-infectious inflammatory condition and initiate empiric corticosteroids, which can exacerbate an occult infection(6). Furthermore, even with negative microbiologic testing, providers often continue empiric antibiotics due to concerns of falsely negative results, a practice that drives emergence of antibiotic resistance and increases risk of *Clostridium difficile* infection(7). In the intensive care unit (ICU), LRTI diagnosis is particularly complex due to a high prevalence of non-infectious inflammatory conditions with overlapping clinical features(8) and a patient demographic that includes severely immunocompromised individuals who may exhibit atypical presentations of pulmonary infections.

Advancements in genome sequencing hold promise for overcoming these diagnostic challenges by affording culture-independent assessment of microbial genomes from microliter volumes of clinical samples(9, 10). Recent work has highlighted the utility of metagenomic next-generation sequencing (mNGS) for rapid and actionable diagnosis of complicated infections(6, 11–13). While these results are encouraging, most mNGS computational pipelines have been developed for analysis of sterile fluids or cultured bacterial isolates and have limited capacity to identify pathogens amidst the complex background of commensal microbiota present in respiratory specimens^13–15^.

Host transcriptional profiling from peripheral blood has emerged as a promising alternative to pathogen-based diagnostics that can distinguish viral from bacterial LRTIs as well as differentiate between patients with acute respiratory infections versus those with non-infectious illnesses(5, 16, 17). This approach, while highly promising, has not been well studied in ICU patients with respiratory failure or in severely immunocompromised subjects. Furthermore, host transcriptional profiling has not yet been coupled with simultaneous detection of pulmonary pathogens(5, 18), which could improve diagnostic accuracy and more precisely inform optimal antimicrobial treatment.

mNGS can extend both host gene expression assays and current microbe-based diagnostics by simultaneously detecting pathogens, the airway microbiome, and transcriptional biomarkers of the host’s immune response. Here we address the need for better LRTI diagnostics by developing an mNGS-based method that integrates host response and unbiased microbe detection. We then evaluate the performance of this approach in a prospective cohort of critically ill patients with acute respiratory failure.

## RESULTS

We prospectively enrolled 92 adults admitted to the ICU with acute respiratory failure and collected tracheal aspirate (TA) samples within 72 hours of intubation (**Table 1**). Patients underwent testing with clinician-ordered standard of care microbiologic diagnostics at the University of California San Francisco Moffitt-Long Hospital, a tertiary care referral center. Subjects with LRTI were identified by two-physician adjudication using United States Centers for Disease Control/National Healthcare Safety Network (CDC/NHSN) surveillance case definitions and retrospective electronic medical record review, with blinding to mNGS results (**Table S2A**)(19). Using this approach, patients were assigned to one of four groups: 1) LRTI defined by both clinical and microbiologic criteria (LRTI^+C+M^, n=26); 2) no evidence of LRTI and a clear alternative explanation for acute respiratory failure (no-LRTI, n=18), 3) LRTI defined by clinical criteria alone with negative conventional microbiologic testing (LRTI^+C^, n=34) and 4) respiratory failure due to unclear cause, infectious or non-infectious (unk-LRTI, n=14).

**Table 1.** Demographics and clinical characteristics of study cohort. Legend: LRTI^+C+M^ = subjects who met both clinical and microbiologic criteria for LRTI. no-LRTI = subjects with a non-infectious etiology of acute respiratory failure. SIRS = systemic inflammatory response syndrome, defined as two or more abnormalities in white blood cell count (>12,000 cells/µL or <4,000 cells/µL), temperature (>38°C or < 36°C), heart rate (>90 beats per minute) or respiratory rate (> 20 breaths per minute). APACHEIII score predicts mortality and disease severity for critically ill patients. Pneumonia severity index score estimates mortality for adult patients with community-acquired pneumonia(68). COPD = chronic obstructive pulmonary disease. ^♦^Chi-squared test. *Wilcoxon rank sum test.

| Patient Characteristics | Cohort Overall |  | LRTI<sup>+C+M</sup> | no-LRTI | p<sup>†</sup> |
| --- | --- | --- | --- | --- | --- |
| Total enrolled | 92 |  | 26 | 18 | - |
| Age, years (mean, range) | 62 (21-85+) |  | 61 | 63 | 0.80 |
| Female gender | 31 | 34% | 6 | 9 | 0.13 |
| Race: |  |  |  |  |  |
| African American | 5 | 5% | 2 | 1 | 0.82 |
| Asian | 26 | 28% | 7 | 5 |  |
| Caucasian | 50 | 46% | 15 | 9 |  |
| Other | 11 | 12% | 2 | 3 |  |
| Hispanic Ethnicity | 8 | 9% | 3 | 1 | 0.88 |
| Comorbidities and Outcomes |  |  | LRTI<sup>+C+M</sup> | no-LRTI | p<sup>†</sup> |
| Bacteremia | 21 | 23% | 6 | 3 | 0.90 |
| Non-pulmonary infections | 29 | 32% | 9 | 4 | 0.58 |
| COPD | 12 | 13% | 3 | 0 | 0.37 |
| Diabetes mellitus | 6 | 7% | 1 | 3 | 0.36 |
| Congestive heart failure | 7 | 8% | 1 | 1 | 1.00 |
| Current smoker | 12 | 13% | 5 | 1 | 0.39 |
| Immune suppression | 41 | 45% | 10 | 9 | 0.65 |
| Solid organ transplantation | 13 | 14% | 1 | 5 | 0.07 |
| Prior antibiotic use | 84 | 91% | 22 | 18 | 0.23 |
| Community acquired pneumonia | 42 | 46% | 18 | - | - |
| Hospital acquired pneumonia | 13 | 14% | 5 | - | - |
| Ventilator associated pneumonia | 3 | 3% | 3 | - | - |
| 30-day mortality | 18 | 20% | 6 | 1 | 0.25 |
| Clinical Metrics |  |  | LRTI<sup>+C+M</sup> | no-LRTI | p<sup>*</sup> |
| Max Temperature °C | 37.8 |  | 38.1 | 38.0 | 0.33 |
| Max WBC count (10<sup>6</sup> cells/μL) | 14.3 |  | 13.8 | 12.8 | 0.58 |
| Max Heart Rate (bpm) | 110 |  | 111 | 107 | 0.50 |
| Max Respiratory Rate (breaths/min) | 36 |  | 35 | 35 | 0.74 |
| SIRS Criteria (mean) | 3 |  | 3 | 3 | 0.54 |
| APACHE III score (mean) | 97 |  | 101 | 94 | 0.62 |
| Pneumonia Severity Index (mean) | 151 |  | 148 | 137 | 0.65 |

From extracted nucleic acid samples, we performed both metagenomic shotgun DNA sequencing (DNA-Seq) as well as RNA sequencing (RNA-Seq). We first developed computational algorithms to sift respiratory pathogens from background commensal flora in an effort to enhance detection of LRTI etiology. To differentiate patients with LRTI from those with non-infectious critical respiratory illnesses, we next developed metrics of LRTI probability based on pathogen, airway microbiome diversity, and host gene expression (**Fig 1**). To assess assay performance, we focused on the most unambiguously LRTI positive and negative subjects (LRTI^+C+M^ and no-LRTI) by randomly dividing them into independent derivation (n=20, used for model training) and validation cohorts (n=24, used for model testing). Each metric (pathogen, microbiome, host) was evaluated independently and then in combination.

**Figure 1.**
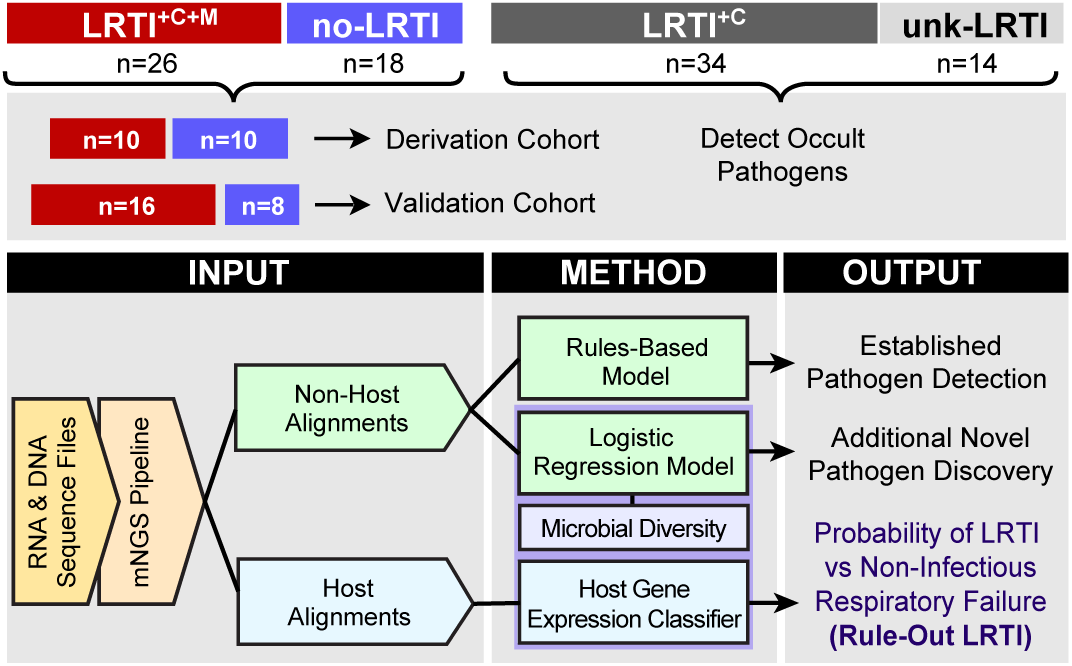
Study overview and novel analysis workflow. Patients with acute respiratory failure were enrolled within 72 hours of ICU admission and TA samples were collected and underwent both RNA sequencing (RNA-Seq) and shotgun DNA sequencing (DNA-Seq). Post-hoc clinical adjudication blinded to mNGS results identified patients with LRTI defined by clinical and microbiologic criteria (LRTI^+C+M^); LRTI defined by clinical criteria only (LRTI^+C^); patients with non-infectious reasons for acute respiratory failure (no-LRTI); and respiratory failure due to unknown cause (unk-LRTI). The LRTI^+C+M^ and no-LRTI groups were divided into derivation and validation cohorts. To detect pathogens and differentiate them from a background of commensal microbiota we developed two models. A rules-based model (RBM) and a logistic regression model (LRM). LRTI probability was next evaluated with i) a pathogen metric, ii) a lung microbiome diversity metric, and iii) a 12-gene host transcriptional classifier. Models were then combined and optimized for LRTI rule-out.

### Pathogen Detection

While many NGS platforms utilize only one nucleic acid type, we combined both RNA and DNA sequencing. This approach allowed for simultaneous host transcriptional profiling, permitted detection of RNA viruses, and enriched for actively transcribing microbes (versus latent or nonviable taxa). In addition, requiring concordant detection of microbes across both nucleic acid types reduced spurious alignments derived from reagent contaminants intrinsic to the library preparations of each nucleic-acid type(20). From each TA sample, we generated a mean of 19.6 and 32.6 million paired-end sequencing reads, from DNA and RNA-Seq respectively, of which the median fraction of microbial reads was 0.04% (IQR 0.01% - 0.16%). Raw reads were analyzed using a rapid computational pipeline that aligns and classifies microbial taxa by nucleotide and peptide translation using the National Center for Biotechnology Information (NCBI) NT and NR databases, respectively(20, 21). RNA-Seq yielded a greater abundance of sequences as compared to DNA-Seq for 78% of identified microbes, with a median of 2.2 times more reads per microbe.

We and others have previously developed NGS methodologies for “sterile site” clinical fluids such as cerebrospinal fluid (CSF)(14, 21, 22). The lung however is not a sterile environment and in fact harbors microbial communities during states of both health and disease(23–26). Asymptomatic carriage of potentially pathogenic organisms is common(27, 28), and only in a subset of cases do these microbes overtake airway microbial communities and precipitate LRTI(29). As such, distinguishing legitimate pathogens from commensal or colonizing microbiota is a central challenge for LRTI diagnostics and adds complexity to the interpretation of metagenomic sequencing data. To this point, while we detected all 38 pathogens identified from clinician-ordered microbiologic tests in the 26 LRTI^+C+M^ patients using mNGS (**Table S3A**), a ten-fold greater number of airway commensals were also identified. The most prevalent microbes in the no-LRTI patient group included well known commensal taxa (**Table S4**). Thus, to distinguish probable pathogens from airway commensals, we developed two complementary algorithms: 1) a rules-based model (RBM) optimized for detecting well-established respiratory pathogens, and 2) a more flexible logistic regression model (LRM) that also permitted novel pathogen detection (**Figure 1**).

The goal of both models was to correctly identify pathogens amidst abundant and heterogeneous populations of commensals. Microbes identified by clinician-ordered diagnostics plus all viruses with established respiratory pathogenicity in the LRTI^+C+M^ group were categorized as pathogens (n=12 in derivation cohort and n=26 in validation cohort, **Table S1**). Any additional microbes identified by mNGS were considered commensals (n = 155 in derivation cohort, n = 174 in validation cohort). We accepted that this “practical” gold standard would provide an attenuated estimate of performance due to the sensitivity limitations of microbial culture in the setting of antibiotic pre-administration(4).

In the RBM, respiratory microbes from each patient were assigned an abundance score based on the sum of log(RNA-Seq) and log(DNA-Seq) genus reads per million reads mapped (rpm) (**Table S3A**). After ranking microbes by this abundance score, the greatest score difference between sequentially ranked microbes was identified and used to distinguish the group of highest-scoring microbes within each patient (**Fig 2A, Fig S1**). These high scoring microbes plus all RNA viruses detected at a conservative threshold of > 0.1 rpm were indexed against an *a priori* developed table of established lower respiratory pathogens derived from landmark surveillance studies and clinical guidelines (**Table S2B**), and if present were identified as putative pathogens by the RBM(4, 30–32).

**Figure 2.**
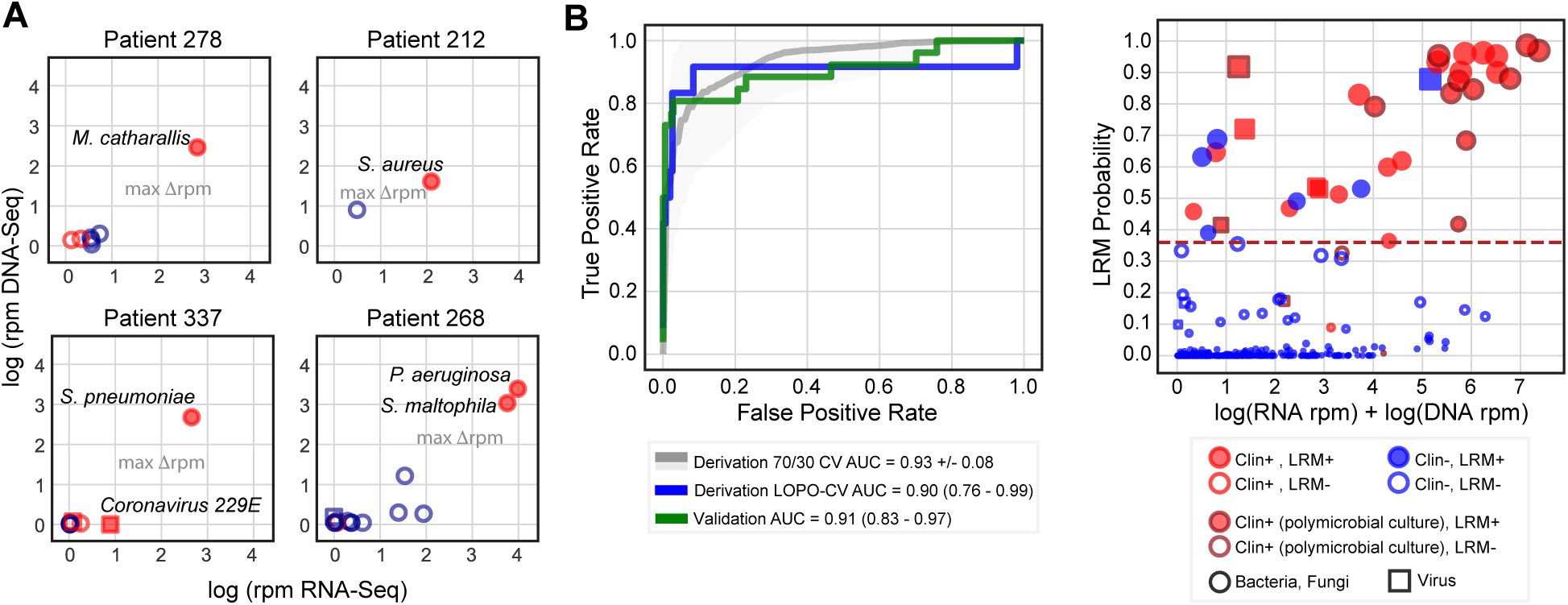
Workflow for distinguishing LRTI pathogens from commensal respiratory microbiota using an algorithmic approach. **A)** Projection of microbial relative abundance in log rpm (reads per million reads sequenced) by RNA sequencing (RNA-Seq, X axis) versus DNA sequencing (DNA-Seq, Y axis) for representative cases. In the LRTI^+C+M^ group, pathogens identified by standard clinical microbiology (filled shapes) had higher overall relative abundance as compared to other taxa detected by sequencing (open shapes). The largest score differential between ranked microbes (max Δrpm) was used as a threshold to identify high-scoring taxa, distinct from the other microbes based on abundance (line with arrows). Red indicates taxa represented in the reference list of established LRTI pathogens. **B)** ROC curve demonstrating logistic regression model (LRM) performance for detecting pathogens versus commensal microbiota in both the derivation and validation cohorts. The grey ROC curve and shaded region indicate results from 1000 rounds of training and testing on randomized sets the derivation cohort. The blue and green lines indicate predictions using leave-one-patient-out cross-validation (LOPO-CV) on the derivation and validation on the validation cohort, respectively. **C)** Microbes predicted by the LRM to represent putative pathogens. The X axis represents combined RNA-Seq and DNA-Seq relative abundance, the Y axis indicates pathogen probability. The dashed line reflects the optimized probability threshold for pathogen assignment. <u>Legend:</u> <u>*Red filled circles*:</u> microbes predicted by LRM to represent putative LRTI pathogens that were also identified by conventional microbiologic tests. *<u>Blue filled circles:</u>* microbes predicted to represent putative LRTI pathogens by LRM only. <u>*Blue open circles*:</u> microbes identified by NGS but not predicted by the LRM to represent putative pathogens. <u>*Red open circles:*</u> microbes identified using NGS and by standard microbiologic testing but not predicted to be putative pathogens. <u>*Dark red outlined circles*:</u> microbes detected as part of a polymicrobial culture.

The RBM achieved an accuracy for pathogen detection of 98.8% and 95.5% in the derivation and validation cohorts, respectively (**Table S3A**). In subjects whose respiratory cultures grew three or more different bacteria, mNGS was able to detect each of the species. In most cases however, their abundance differed by several hundred-fold, which confounded detection of the lower abundance taxa (**Table S3A**). Given the unclear significance of single species in such polymicrobial cases with respect to pathogenicity(33), we performed a secondary analysis in which only the most abundant microbe was considered a pathogen, and this approach yielded an accuracy of 98.4%.

While the RBM performed well for identifying microbes with established pulmonary pathogenicity, we recognized the need to also detect novel or atypical species. We thus employed machine learning to distinguish respiratory pathogens from commensals using a logistic regression model (LRM) trained on microbes detected in the derivation cohort patients (n=20) using the predictor variables of: RNA-Seq rpm, DNA-Seq rpm, rank by RNA-Seq rpm, established LRTI pathogen (yes/no), and virus (yes/no). These features were selected to preferentially favor highly abundant organisms with established pathogenicity in the lung, but still permit detection of uncommon taxa that could represent putative pathogens.

To evaluate LRM performance in the derivation cohort, we performed leave-one-patient-out cross validation, in which all microbes from a single patient were held out in each round of cross-validation. This yielded an AUC = 0.90 (95% CI = 0.76 – 0.99). A final model was trained on all microbes from derivation cohort patients, and this achieved an AUC = 0.91 (95% CI = 0.83– 0.97) for pathogen identification in the validation cohort (**Fig 2B, Table S3A, S3B**). At an optimized probability threshold of 0.36 (**Methods**), this translated to an accuracy of 96.4% and 95.5% in the derivation and validation cohorts, respectively. As with the RBM, LRM performance suffered in polymicrobial culture cases with species that differed by several magnitudes in abundance when assessed by mNGS. As such, when only the most abundant microbe identified by clinical microbiologic diagnostics per LRTI^+C+M^ patient was considered as the etiologic pathogen, the AUC increased to 0.997 (95% CI = 0.99 – 1.00) in the validation cohort.

Combining the RBM and LRM identified more putative pathogens than either model alone and revealed a potential LRTI etiology in 62% (n=21) of the LRTI^+C^ patients with clinically adjudicated LRTI but negative microbiologic testing (**Figs 3, S2, Table S3A**). Compared to clinician-ordered diagnostics, this permitted a microbiologic diagnosis in a greater number of LRTI-positive subjects (78% vs 43%, p < 1.00 x 10^−4^ by McNemar’s test, **Fig 3**). Putative new pathogens in a representative subset of the LRTI^+C^ group patients (n=11, 32%) were orthogonally confirmed by clinical multiplex respiratory virus PCR, influenza C PCR(34) or by 16S bacterial rRNA gene sequencing (**Table S3A**).

**Figure 3.**
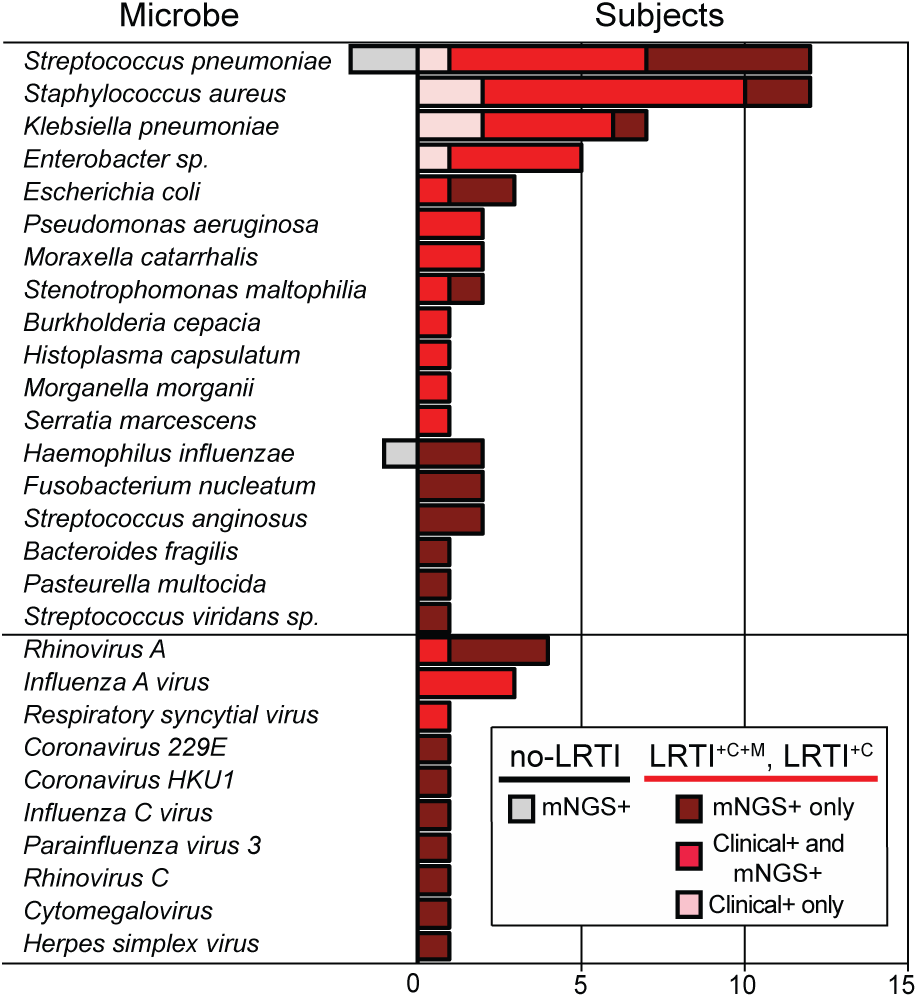
Distribution of respiratory pathogens identified in patients using clinician-ordered diagnostics versus mNGS. Number of subjects in whom each respiratory microbe was detected. All microbes detected by clinician-ordered diagnostics were detected by mNGS, however pink bars indicate microbes misclassified as negative by either the rules-based or logistic regression models. Notably, all microbes identified by clinician-ordered diagnostics and misclassified by either the rules-based or logistic regression models (pink bars) were found in polymicrobial cultures, highlighting the presence of dominant pathogens by NGS that are not captured in the polymicrobial culture results. Red bars indicate microbes detected by clinician-ordered diagnostics and also predicted as pathogens by either the rules based or logistic regression models. More detail on which model identified each microbe can be found in Figure S2. Dark red bars (LRTI^+C+M^ and LRTI^+C^ subjects) and grey bars (no-LRTI subjects) indicate number of cases with microbes detected only by mNGS.

Putative pathogens identified in the unk-LRTI group (n=6, 42%) may have represented atypically presenting respiratory infections or incidental carriage in the respiratory tract (**Fig S2, Table S3A**). Microbes identified in the no-LRTI group (n=3, 17%) were present at lower abundance compared to microbes in LRTI^+C+M^ subjects (p < 0.01 by Wilcoxon rank sum), LRTI^+C^ (p < 0.01) and unk-LRTI subjects (p = 0.02), and included contextual pathogens such as *S. pneumoniae and H. influenzae* that colonize the airways of 20-50% of healthy individuals(33, 35, 36). Together, these findings highlighted the reality of asymptotic carriage of potentially pathogenic species, emphasizing the need to contextualize microbial detection with respect to other key elements of an airway infection, in particular the airway microbiome and the host’s immune response(27, 37). We thus undertook further analytical development to predict LRTI status by calculating combined metrics based on pathogen, microbiome and host transcriptional response.

### LRTI Prediction Based on Pathogen

We recognized that the highest per-patient LRM pathogen versus commensal probability value differed significantly between LRTI^+C+M^ and no-LRTI subjects (p = 3.8 x 10^−4^ by Wilcoxon rank sum). As such, we hypothesized that this value might have utility not only for pathogen versus commensal prediction, but also for LRTI prediction in general. Testing this idea, we found that the maximum per patient LRM probability value predicted LRTI status with an AUC of 0.97 (95% CI = 0.90 - 1.00) in the derivation cohort and 0.96 (95% CI = 0.86 - 1.00) in the validation cohort (**Fig S3**).

### LRTI Prediction Based on Lung Microbiome Diversity

Several studies have demonstrated reduced diversity of the airway microbiome in the setting of LRTI(20, 38–40). We measured intra-patient (alpha) diversity of airway genera using the Shannon Diversity Index (SDI) and found that LRTI^+C+M^ subjects had significantly lower SDI compared to no-LRTI subjects when assessed by both RNA-Seq (**Fig 4A,** p = 1.3 x 10^−4^) and DNA-Seq (**Fig S4A**, p = 8.9 x 10^−3^) (**Table S5**). We next examined inter-patient (beta) diversity(41) using the Bray-Curtis Index(42) and found that this also differed between LRTI^+C+M^ and no-LRTI subjects, with assessment by RNA-Seq again yielding a more significant difference versus DNA-Seq (p = 5 x 10^−3^ versus p = 9 x 10^−3^ by PERMANOVA, respectively, **Figs 4B and S4B**). We then tested whether diversity alone might predict LRTI and found that RNA-Seq SDI differentiated LRTI^+C+M^ from no-LRTI subjects with an AUC of 0.96 (95% CI = 0.89 - 1.00) in the derivation cohort and an AUC of 0.80 (95% CI = 0.63 – 0.96) in the validation cohort (**Fig 4C**). DNA-Seq SDI did not perform as well, with AUCs of 0.84 (95% CI = 0.66 – 1.00) and 0.53 (95% CI = 0.25 – 0.80) in the derivation and validation cohorts, respectively (**Fig S4C**). These findings suggested that genus diversity assessed by RNA-Seq was a useful, albeit imperfect, biomarker of LRTI.

**Figure 4.**
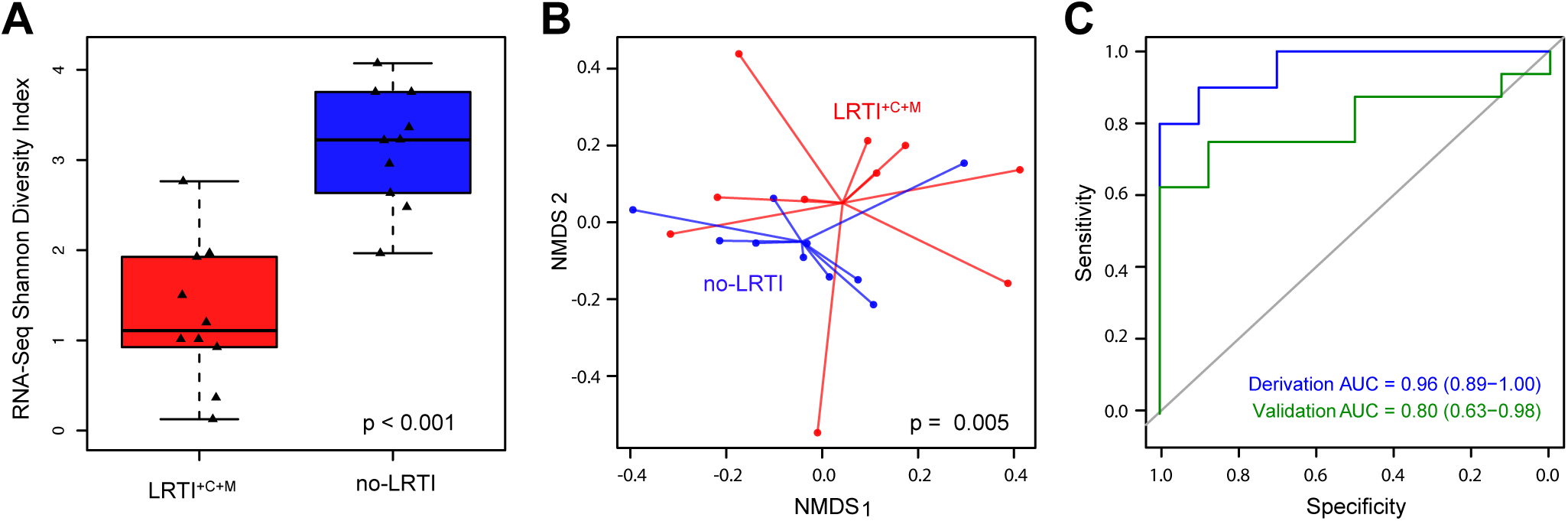
Diversity of the transcriptionally active lung microbiome in patients with LRTI (LRTI^+C+M^) versus non-infectious respiratory illnesses (no-LRTI). **A)** Box plots of Shannon Diversity Index (SDI) of the lung microbiome assessed by RNA-Seq at the genus level (in the derivation cohort) differed between LRTI^+C+M^ from no-LRTI groups. **B)** Beta diversity assessed by PERMANOVA on Bray-Curtis dissimilarity values in the derivation cohort differed between LRTI^+C+M^ and no-LRTI groups. **C)** ROC curve demonstrating performance of SDI to distinguish LRTI^+C+M^ from no-LRTI groups.

### LRTI Prediction Based on Host Response

In the setting of critical illness, systemic inflammatory responses due to diverse physiologic processes can make true LRTI clinically indistinguishable from non-infectious respiratory failure or severe extra-pulmonary infection. Consistent with this, we found that the systemic inflammatory response syndrome (SIRS) criteria (temperature, white blood cell count, heart rate, respiratory rate) had limited utility for LRTI detection despite being widely used for infection assessment (**Table S1**). We thus hypothesized that transcriptional profiling, which has emerged as a promising and accurate host-based approach for assessing infection, might provide diagnostic insight in settings when clinical rules are uninformative(5, 16, 43).

As such, we examined differential gene expression between LRTI^+C+M^ and no-LRTI subjects in the derivation cohort to define a host transcriptional signature of LRTI in patients with critical illness. Using a false discovery rate (FDR) of < 0.05, we identified a total of 882 differentially expressed genes, 414 of which were upregulated in LRTI^+C+M^ subjects (**Fig S6, Table S6A**). Gene set enrichment analysis(44) identified upregulation of pathways related to innate immune responses, NF-κβ signaling, cytokine production, and the type I interferon response in LRTI^+C+M^ subjects. In comparison, gene expression pathways in the no-LRTI group were enriched for oxidative stress responses and MHC class II receptor signaling (**Table S6B**). A sub-analysis (**Supplemental Methods**) evaluating differences between viral and bacterial infections in known LRTI^+C+M^ patients identified four differentially expressed genes (RSAD2, OAS3, CXCL2, DUSP2). Genes upregulated in viral cases (RSAD2, OAS3) were related to the type-1 interferon and anti-viral responses, reflecting biologically relevant differences in host response indicative of pathogen type, despite a relatively limited sample size within a heterogeneous cohort and high proportion of immune compromising conditions in the majority patients with detected viruses.

We next sought to construct an airway-specific host transcriptional classifier that could differentiate LRTI^+C+M^ patients from no-LRTI subjects by employing machine-learning (**Methods**). Elastic net regularized regression in the derivation cohort identified a 12-gene classifier that was then used to score patients based on a weighted sum of scaled expression values (**Table S7A, Fig 5A, B**). We found that predictive classifier genes upregulated in LRTI^+C+M^ patients compared to no-LRTI patients included *NFAT-5*, which plays a role in T cell function and inducible gene transcription during immune responses(45); *ZC3H11A*, which encodes a zinc-finger protein involved in the regulation of cytokine production and immune cell activation(46) and *PRRC2C*, which functions in RNA binding and may play a role in hematopoietic progenitor cell differentiation in response to infection(47). Genes upregulated in no-LRTI patients compared to LRTI^+C+M^ patients included: *CD36*, which encodes a macrophage phagocytic receptor involved in scavenging dying/dead cells and oxidized lipids(48, 49); *BLVRB*, which is involved in oxidative stress responses(50), *EDF1*, which contributes to the regulation of nitric oxide release in endothelial cells(51) and *ENG*, an integral membrane glycoprotein receptor that may modulate inflammation and angiogenesis(52).

**Figure 5.**
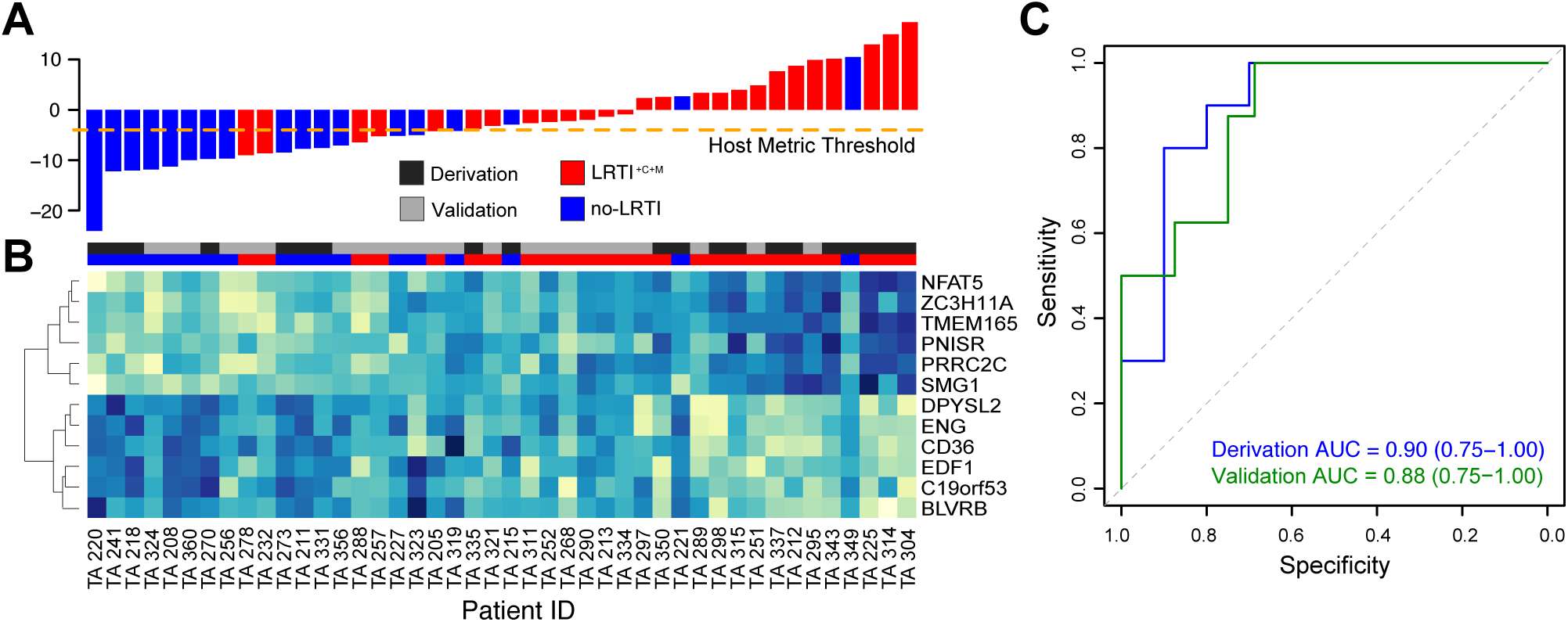
Host transcriptional profiling distinguishes patients with acute LRTI (LRTI^+C+M^) from those with non-infectious acute respiratory illness (no-LRTI). **A)** Host classifier scores for all patients in the derivation and validation cohorts, each bar indicates a patient score and is colored as follows: LRTI^+C+M^ = red, no-LRTI = blue. Orange dotted line indicates the host classifier threshold (score = −4) that achieved 100% sensitivity in the training set and was used to classify the test set samples. **B)** Normalized expression levels, arranged by unsupervised hierarchical clustering, reflect over-expression (blue) or under-expression (turquoise) of classifier genes (rows) for each patient (columns). 12 genes were identified as predictive in the derivation cohort and subsequently applied to predict LRTI status in the validation cohort. Column colors above the heatmap indicate whether a patient belonged to the derivation cohort (dark grey) or validation cohort (light grey) and whether they were adjudicated to have LRTI^+C+M^ (red) or no-LRTI (blue)**. C)** ROC curves demonstrating host classifier performance for derivation (blue) and validation (green) cohorts.

Classifier performance assessed by leave-one-out cross-validation demonstrated an AUC of 0.90 (95% CI 0.75 – 1.00) in the derivation cohort and an AUC of 0.88 (95% CI 0.75 – 1.00) in the validation cohort (**Fig 5C**). Covariates for immune suppression, concurrent non-pulmonary infection, antibiotic use, age, and gender were iteratively incorporated into the regression model but none were significant enough to be maintained when sparsity was added by elastic net (**Table S7B**). We tested whether differences in host gene expression could be attributed to enrichment of specific cell types using CIBERSORT(53) (**Table S7C**) and found that only M2 Macrophages were enriched in the no-LRTI group (p = 0.03 by Wilcox Rank Sum).

Finally, given our modest sample size, we tested the statistical power of our host classifier by computing learning curves (**Methods**). We observed that even with subsampling, the 12 classifier genes were continually represented. While the derivation cohort sample size approached the limit required for robust performance assessment, the analysis suggested that additional patients might lead to further improvement (**Figure S6A**). A similar analysis for the pathogen versus commensal LRM indicated that performance metrics had converged with the given microbial sample size, indicating robust performance assessment and sufficient training data (**Figure S6B**).

### Evaluation of a Combined LRTI Metric

Given the relative success of each independent metric (pathogen, microbiome and host) for discerning presence of infection, we asked whether combining them could enhance LRTI detection. We recognized the potential of mNGS to empower a data-driven assessment of a patient’s LRTI status during the critical timeframe following ICU admission. As such, we developed a readily interpretable compilation of host and pathogen mNGS metrics in a *rule-out model* designed to maximize LRTI diagnostic sensitivity. This process, which involved optimizing intra-metric LRTI positivity thresholds in the derivation cohort and calling positivity based on either the host <u>or</u> pathogen scores (**Methods**), achieved a sensitivity and specificity of 100% and 87.5%, respectively in the validation cohort, equating to a negative predictive value of 100% (**Fig 6B)**. Despite the limitations of a small cohort, we investigated the potential utility of the *rule-out model* for curbing broad-spectrum antibiotic overuse in the ICU by performing a theoretical calculation in the no-LRTI group to estimate the potential impact of mNGS result availability at 48 hours post-enrollment. This estimate suggested that a significant reduction in unnecessary empiric antibiotic use could have been possible (78 versus 50 days of therapy, p = 0.03, **Supplemental Methods**).

**Figure 6.**
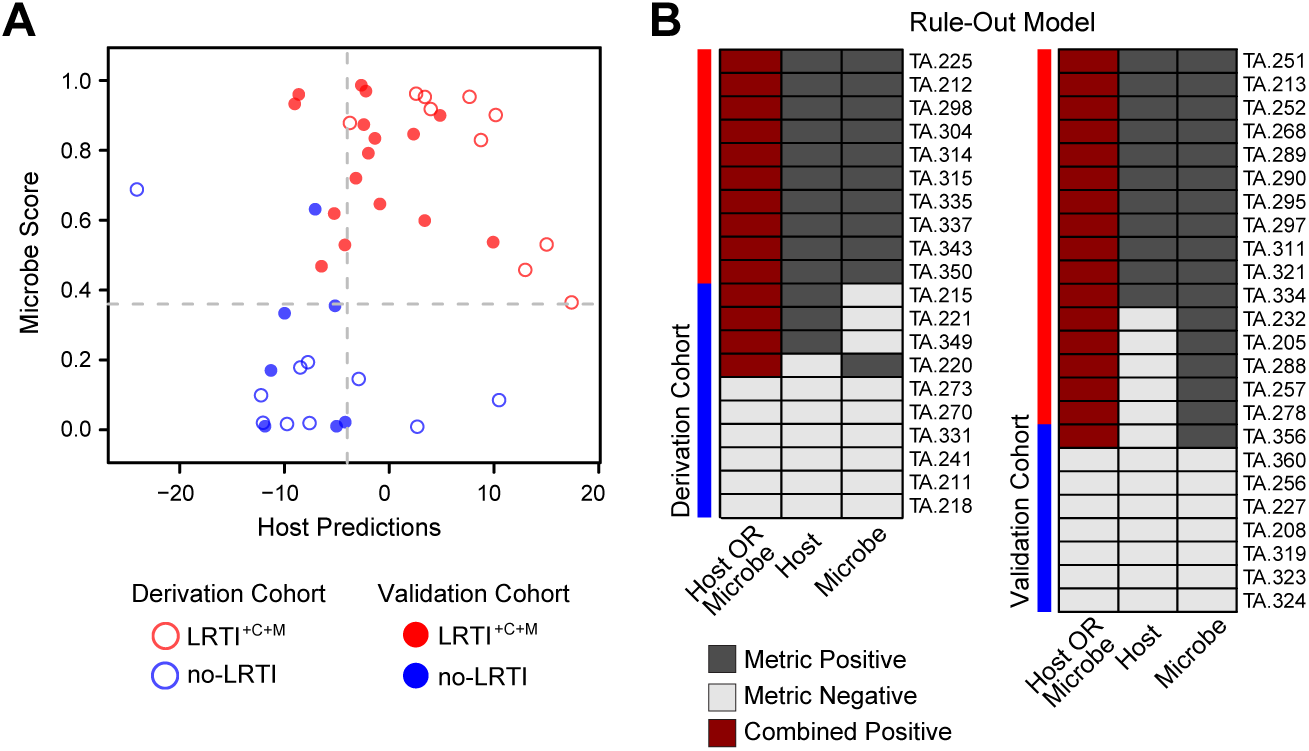
Combined LRTI prediction metric integrating pathogen detection and host gene expression. **A)** Scores per patient for each of the two components of this *LRTI rule-out model* are projected into a scatterplot (X axis represents the host metric, Y axis represents the microbe score). The thresholds optimized for sensitivity in the derivation cohort are indicated in grey dashed line. Each point represents one patient – those that were in the derivation cohort have no fill and those that were in the validation cohort are filled. Red indicates LRTI^+C+M^ and blue indicates no-LRTI subjects. **B)** LRTI rule-out model results for each patient are shown for both the derivation and validation cohorts, with study subjects shown in rows and metrics in columns. Dark grey indicates a metric exceeded the optimized LRTI threshold, light grey indicates it did not. Dark red indicates the subject was positive for both pathogen + host metrics, and thus was classified as having LRTI. White indicates missing data.

## DISCUSSION

Of all infectious disease categories, LRTIs impart the greatest mortality both worldwide and in the United States(1). Contributing to this is the rising rate of treatment failure due to antibiotic resistance(54) and the limited performance of existing diagnostics for identifying respiratory pathogens(4, 55). In this prospective cohort study, we describe the use of unbiased mNGS for respiratory infectious disease diagnosis in the ICU. We develop methods that advance pathogen-based genomic diagnostics as well as existing host transcriptional classifier platforms by simultaneously assessing respiratory pathogens, the airway microbiome, and the host transcriptome in a single test to predict LRTI and identify disease etiology. We find that host/pathogen mNGS accurately detects LRTI in patients with acute respiratory failure and can provide a microbiologic diagnosis in cases due to unknown etiology.

Host transcriptional profiling has gained attention as a promising approach to LRTI diagnosis(56, 57) but is understudied in critically ill and immunocompromised patients, who may be the most likely to benefit from this technology. We addressed this gap by interrogating airway gene expression in a critically ill cohort with 45% immunocompromised patients to develop an accurate host transcriptional classifier. Unlike existing classifiers, host-microbe mNGS offers the advantage of simultaneous species-level microbial identification.

The role of commensal lung microbiota in health and disease is an area of active investigation. We corroborated prior findings demonstrating microbiome differences between subjects with respiratory infections and those with non-infectious airway disease(20, 38). More specifically, we found that LRTI was associated with reduced intra-patient alpha diversity of the airway microbiome and that collectively, patients with LRTI differed significantly from those without in terms of beta diversity and microbial burden. This diversity difference was more pronounced when assessed by RNA-Seq, potentially due to inclusion of RNA viruses and transcripts from actively replicating pathogens in infected patients. As a biomarker, RNA-Seq SDI had moderate utility for predicting LRTI; however, it did not enhance performance in combination with the other metrics, perhaps due to negative correlation with microbe score (r = −0.84 in the derivation cohort).

Discriminating respiratory pathogens from background commensal microbiota is a key challenge for LRTI diagnostics and is particularly relevant for sensitive molecular assays(58). We directly addressed this by developing two complementary algorithms (RBM, LRM) that parsed putative pathogens from airway commensals. When combined, these models enabled a microbiologic diagnosis in significantly more patients with LRTI compared to clinician-ordered diagnostics. The fact that the *a. priori* selected model features successfully differentiated pathogens from commensals validated the underlying model assumptions related to pathogen dominance resulting in disruption of alpha diversity. Notably, both models also proved useful despite widespread antibiotic use prior to airway sampling (90% of subjects), a practice that occurs commonly and that can sterilize microbial cultures(59).

The capacity for mNGS to detect pathogens unidentifiable by standard clinical diagnostics was highlighted in several cases, including that of subject 254, who developed rapidly worsening respiratory failure and fever during a prolonged post-surgical admission. He was treated empirically for hospital acquired pneumonia with linezolid, aztreonam and metronidazole. Lower respiratory cultures returned negative, but mNGS identified influenza C, which is not available on most clinical multiplex viral PCR assays. Notably, 12% of subjects were found to have undetected and potentially transmissible respiratory viruses despite strict precautionary respiratory contact policies at the study site, a finding that suggests the potential value of mNGS for hospital infection control. Several cases also highlighted the potential for mNGS to enhance antibiotic stewardship, and we estimated that theoretical implementation of the *rule-out model* within 48 hours could have reduced antibiotic days of therapy by 36% in the no-LRTI validation cohort patients.

Since at the time of ICU admission it is often difficult to distinguish infectious from non-infectious acute respiratory disease, a theoretical workflow for host/microbe mNGS could involve first employing the *rule-out model* to asses LRTI probability and complement clinical decision making regarding discontinuation of empiric antimicrobials. In cases where LRTI was ultimately suspected, a microbiologic diagnosis could then be obtained using a combination of the RBM and LRM to accurately screen for both well-established and uncommon respiratory pathogens. A principal advantage of mNGS is that all potential infectious agents can be simultaneously assessed, which avoids the need for ordering multiple individual tests for each different pathogen of concern. Future studies in a larger validation cohort can help optimize host and microbe LRTI rule-out thresholds and further assess test performance prior to deployment in a clinical setting.

Some limitations of host/microbe mNGS were apparent and included false-positive detection of pathobionts such as *H. influenzae* and *S. pneumoniae* in the no-LRTI group, and false positivity of the host-response metric in subjects including patient 349, who was diagnosed with alpha-1 antitrypsin deficiency-associated pulmonary disease. The relatively small sample size of our derivation and validation cohorts increased the potential for data overfitting and was a limitation of our study. Learning curve estimates, however, indicated that the sample size was optimal for pathogen versus commensal prediction, and adequate for the host classifier, consistent with the estimate from an established sample size prediction tool for high dimensional classifiers (60) (**Supplemental Methods**). Nonetheless, a larger cohort will be necessary to improve the robustness of model performance estimates and better assess synergy resulting from combining host and microbial metrics.

Strengths of this study include an innovative bioinformatics approach, detailed patient phenotyping, and a study population reflective of the true heterogeneity of ICU patients, including severely immunocompromised subjects and patients receiving broad spectrum antibiotics. Future studies in a larger cohort can further validate these findings, strengthen the utility of these models, and assess the impact of mNGS on clinical outcomes. In summary, we report a multifaceted approach to LRTI diagnosis that is the first to integrate three central elements of airway infections: the pathogen, airway microbiome and host’s response.

## METHODS

### Study Design and Subjects

This prospective observational study evaluated adults with acute respiratory failure requiring mechanical ventilation who were admitted to the University of California San Francisco (UCSF) Moffitt-Long Hospital ICUs. Subjects were enrolled sequentially between 7/25/13 and 10/17/17 within the first 72 hours of intubation for respiratory failure. The UCSF Institutional Review Board approved an initial waiver consent for obtaining excess respiratory fluid, blood and urine samples, and informed consent was subsequently obtained from patients or their surrogates for continued study participation. For patients whose surrogates provided informed consent, follow-up consent was then obtained if patients survived their acute illness and regained the ability to consent. For subjects who died prior to consent being obtained, a full waiver of consent was approved. For all surviving subjects, if consent was not eventually obtained from either patient or surrogate, all specimens were discarded.

### Clinical Microbiologic Testing

During the period of study enrollment, subjects received standard of care microbiologic testing ordered by the treating clinicians. Respiratory testing from TA, bronchial alveolar lavage (BAL) or mini-BAL included: bacterial and fungal stains and semi-quantitative cultures (n=90); AFB stains and cultures (n=8); 12-target clinical multiplex PCR (Luminex) for influenza A/B, respiratory syncytial virus (RSV), human metapneumovirus (HMPV), human rhinovirus (HRV), adenovirus (ADV) and parainfluenza viruses (PIV) 1-4 (n=23); *Legionella* culture (n=1); *Legionella pneumophila* PCR (n=4), cytomegalovirus (CMV) culture (n=4) and cytology for *Pneumocystis jiroveccii* (n=4). Other microbiologic testing included blood culture (n=89); urine culture (n=87); serum cryptococcal antigen (n=4); serum galactomannan (n=1); serum β-D-glucan (n=1).

### Definitions and Clinical Adjudication of LRTI

Because admission diagnoses made by treating clinicians at the time of study enrollment were by necessity based on incomplete clinical, microbiologic and treatment outcome information, a *post-hoc* adjudication approach was carried out to enhance accuracy of LRTI diagnosis. For this, two attending physicians (one from infectious disease [CL], one from pulmonary medicine [FM]) blinded to mNGS results, retrospectively reviewed each patient’s medical record following hospital discharge or death to determine if they met the United States Centers for Disease Control/National Healthcare Safety Network (CDC/NHSN) surveillance definition of pneumonia, with respect to clinical and/or microbiologic criteria (**Table S1**)(19). Chart review consisted of in-depth analysis of complete patient histories, including laboratory and radiographic results, inpatient notes, and post-discharge clinic notes. Using this approach, subjects were assigned to one of four groups, consistent with a recently described approach(56): 1) LRTI defined by both clinical and laboratory criteria; 2) no evidence of respiratory infection and with a clear alternative explanation for respiratory failure (no-LRTI); 3) LRTI defined by clinical criteria only (LRTI^+C^) and unknown, LRTI possible (unk-LRT). A determination of noninfectious etiology was made only if an alternative diagnosis could be established and results of standard clinical microbiological testing for LRTI were negative.

### Host/Microbe Metagenomic Next-Generation Sequencing

Excess TA was collected on ice, mixed 1:1 with DNA/RNA Shield (Zymo) and frozen at - 80°C. RNA and DNA were extracted from 300µl of patient TA using bead-based lysis and the Allprep DNA/RNA kit (Qiagen). RNA was reverse transcribed to generate complementary DNA (cDNA) and used to construct sequencing libraries using the NEBNext Ultra II Library Prep Kit (New England Biolabs). DNA underwent adapter addition and barcoding using the Nextera library preparation kit (Illumina) as previously described(20). Depletion of abundant sequences by hybridization (DASH) was employed to selectively deplete human mitochondrial cDNA, thus enriching for both microbial and human protein coding transcripts(61). The final RNA-Seq and DNA sequencing (DNA-Seq) libraries underwent 125 nucleotide paired-end Illumina sequencing on a HiSeq 4000.

### Pathogen Detection Bioinformatics

Detection of host transcripts and airway microbes leveraged a custom bioinformatics pipeline(20) that incorporated quality filtering using PRICESeqfilter^24^ and alignment against the human genome (NCBI GRC h38) using the STAR(63) aligner to extract genecounts. To capture respiratory pathogens, additional filtering to remove Pan troglodytes (UCSC PanTro4) using STAR and removal of non-fungal eukaryotes, cloning vectors and phiX phage was performed using Bowtie2(64). The identities of the remaining microbial reads were determined by querying the NCBI nucleotide (NT) and non-redundant protein (NR) databases using GSNAP-L and RAPSEARCH2, respectively.

Microbial alignments detected by RNA-Seq and DNA-Seq were aggregated to the genus-level and independently evaluated to determine genus alpha diversity as described below. The sequencing reads comprising each genus were then evaluated for taxonomic assignment at the species level based on species relative abundance as previously described(20). For each patient, the top 15 most abundant taxa by RNA rpm were identified and evaluated under the requirement that all bacteria, fungi, and DNA viruses had concordant detection of their genomes by DNA-Seq and concordant alignments in NR and NT. RNA viruses did not require concordant DNA-Seq reads. (**Figure 2 and Table S3A**). To differentiate putative pathogens from commensal microbiota, we developed rules-based (RBM) and logistic regression (LRM) methods and benchmarked each on sequencing data from LRTI^+C+M^ and no-LRTI subjects.

### Statistical Analysis

Statistical significance was defined as p less than 0.05, using two-tailed tests of hypotheses. Categorical data were analyzed by chi-squared test and nonparametric continuous variables were analyzed by Wilcoxon rank-sum. For statistical validation in the pathogen versus commensal and LRTI prediction metrics, 10 LRTI^+C+M^ and 10 no-LRTI cases were randomly assigned to create a derivation cohort. Model performance was assessed in an independent validation cohort consisting of 16 LRTI^+C+M^ and 8 no-LRTI cases.

### Pathogen versus Commensal Models

We found that all clinically-confirmed LRTI pathogens were present within the top 15 most abundant microbes by RNA-Seq rpm, which on average represented 99% of reads across all samples. We thus limited analysis to the 15 most abundant NGS-detected genera in each sample. For both models, microbes identified using clinician-ordered diagnostics and all viruses with established respiratory pathogenicity in the derivation cohort subjects were considered “pathogens.” Any additional microbes identified by mNGS in these subjects were considered “commensals”. This equated to 12 “pathogens” and 155 “commensals” in the 20 derivation cohort patients and 26 “pathogens” and 174 “commensals” in the 24 validation cohort patients.

#### Rules-Based Model (RBM)

This model leveraged previous findings demonstrating that microbial communities in patients with LRTI are characterized by one or more dominant pathogens present in high abundance(20, 40). Using either RNA-Seq rpm alone (RNA-viruses) or the combination of RNA-Seq and DNA-Seq rpm (all others), this model identified the subset of microbes with the greatest relative abundance in each sample, which consisted of single microbes in cases of a dominant pathogen and also identified co-infections where several microbes were present within a similar range. All viruses detected by RNA-Seq at > 0.1 rpm and present within the *a. priori –* developed reference index of established respiratory pathogens were considered putative pathogens in the model. The remaining taxa (bacteria, fungi, and DNA viruses) were then aggregated at the genus level, assigned an abundance score based on (log(RNA-Seq rpm) + log(DNA-Seq rpm)), and sorted in descending order by this score. The greatest change in abundance score between sequentially ranked microbes was identified, and all genera with an abundance score greater than this threshold were then evaluated at the species level, by identifying the most abundant species within each genus. If the species was present within the *a. priori –* developed reference index of established respiratory pathogens, it was selected as a putative pathogen by the model (**Fig 2**).

#### Logistic Regression Model (LRM)

This model employed the Python (v 3.6.1) sklearn (v 0.18.1) package to train on distinguishing between “pathogen” and “commensals” using the following five input features: log(RNA-Seq rpm), log(DNA-Seq rpm), per-patient RNA-Seq abundance rank, and two binary variables indicating whether the microbe could be identified in the established index of respiratory pathogens or was a virus. These features were selected in alignment with the observation that the pathogens identified in the LRTI^+C+M^ group were more abundant and within the top-ranked microbes. Moreover, the individual features were significantly different between the pathogens and commensals: (RNA-Seq rpm p = 2.44×10^−4^, DNA-Seq rpm p = 3.55 x 10^−3^, scoring rank p = 3.51×10^−6^). Model performance was estimated in the derivation and validation cohorts and learning curves were computed (**Supplemental Methods**). For identification of etiologic pathogens reported (**Fig 3, Table S3A, Table S2**) the threshold of 0.36 was used for consistency between the LRM for pathogen identification and LRTI detection.

### LRTI Prediction Based on Pathogen

Outside of identifying putative LRTI pathogens, we evaluated whether LRM microbial score alone could be used to classify subjects as LRTI positive or LRTI negative. To do so, we used the top LRM-derived pathogen probability score per patient and evaluated the performance of this value alone to predict likelihood of infection in the LRTI^+C+M^ versus no-LRTI subjects.

### Lung Microbiome Diversity Analysis

Alpha diversity of the respiratory microbiome for each subject was assessed by Shannon Diversity Index (SDI) and Simpson Diversity Index at the genus level using NT rpm and the Vegan (version 2.4.4)(65) package in R (version 3.4.0)(66). Richness (total number of genera) and Burden (combined rpm of all genera) were also evaluated. Viral, bacterial, and fungal microbes were included in all diversity analyses, computed independently for RNA-and DNA-Seq samples without requiring that taxa be concordant on both nucleic acids. Diversity values were then compared between patients with clinically adjudicated LRTI (LRTI^+C+M^) and those with respiratory failure due to non-infectious causes (no-LRTI) using the nonparametric Wilcoxon Rank Sum test. Evaluation of alpha diversity for prediction LRTI status was performed using the SDI value. Beta diversity was evaluated using the Bray-Curtis dissimilarity metric calculated at the genus level using NT rpm and the Vegan package in R. Statistical significance of the beta diversity between LRTI^+C+M^ and no-LRTI patients was assessed using permutation analysis of variance (PERMANOVA, 999 permutations) and the results were visualized using Non-metric Multidimensional Scaling (NMDS).

### Host Gene Expression Analysis

Following quality filtration with PRICESeqfilter(62), RNA transcripts were aligned to the ENSEMBL CRCh38 human genome build using STAR. Subsequently, genes were filtered to include only protein-coding genes that were expressed in at least 50% of patients. All samples used for host transcriptome analysis (both derivation and validation sets) ultimately included more than 95,000 protein-coding genes with an average of 734,844 transcripts per patient.

### Differential Expression Analysis

Gene count data were analyzed using the Bioconductor package DESeq2 (v 1.16.1)(67) in R statistical programming environment. To avoid batch-related confounding and class imbalance, we limited our differential expression analysis to the derivation cohort of 10 LRTI^+C+M^ and 10 no-LRTI samples, sequenced in the same batch. Differentially expressed genes with FDR < .05 were used as input to ToppGene(44) to evaluate for functional pathway enrichment.

### Host Gene Expression Classifier for LRTI Prediction

The derivation cohort was independently normalized using DESeq2 and log-transformed. The values for each gene in the derivation cohort were then scaled and centered by z-score. A classifier was built using the elastic net regularized regression model implementation from the glmnet package (version 2.0.13) in the R Statistical Programming Language (version 3.4.0). Regularization parameter alpha = .5 was selected using leave-one-out cross-validation and optimizing for AUC. To account for heterogeneity in the cohort, the model included covariates of concurrent bloodstream infection, immunosuppression, and gender. No significant difference was seen in these parameters between LRTI^+C+M^ and no-LRTI (**Table S7B**). These covariates were reduced to zero in the model fitting stage. Genes with non-zero weights were used for classification. To obtain a single-value score for each patient, genes selected by the elastic net were evaluated for their correlation with each of the two groups. Genes for which the mean expression was greater in the LRTI^+C+M^ were assigned a weight of 1, and those with mean expression greater in no-LRTI were assigned a weight of −1. The normalized, scaled, expression values for each patient were multiplied by the weight vector and summed across all genes. The total sum was used as a representative score and the AUC was calculated. Given the importance of sensitivity in the context of diagnostics, the threshold selected for analysis of the test cohort and combined metrics (scores = −4) was chosen as the threshold which provided 100% sensitivity in the derivation cohort. The host gene expression classifier was then validated on the validation set and learning curves were used to estimate the reliability of the performance metrics (**Supplemental Methods**).

### Classifier Combination

To generate a readily interpretable compilation of host and microbial mNGS metrics that could enable a data-driven assessment of LRTI, the *rule-out* model was developed. In the *rule*-*out model*, we identified score thresholds from the pathogen and host metrics required to achieve 100% sensitivity in the derivation cohort (pathogen > 0.36, and host > −4) and applied these to the validation cohort to predict LRTI using the following combinatorial rule: *LRTI* = (*HOST*)*positive OR* (*Microbe*)*positive*.

### Identification and Mitigation of Environmental Contaminants

To minimize inaccurate taxonomic assignments due to environmental contaminants, we processed negative water controls with each group of samples that underwent nucleic acid extraction, and included these, as well as positive control clinical samples, with each sequencing run. We directly subtracted alignments to those taxa in water control samples detected by both RNA-Seq and DNA-Seq analyses (**Table S8**) from the raw rpm values in all samples. To account for selective amplification bias of contaminants in water controls resulting from PCR amplification of metagenomic libraries to a fixed standard concentration across all samples, prior to direct subtraction we scaled taxa rpms in the water controls to the median percent microbial reads present across all samples (0.04%). In addition, we confirmed reproducibility of results by sequencing 10% of samples in triplicate and evaluated discrepancies between mNGS and standard diagnostics in a random subset of LRTI^+C^ patients using clinically validated 16S bacterial rRNA gene sequencing and/or viral PCR testing, as described above.

### Data Availability

Raw microbial sequences are available via SRA BioProject Accession ID SUB3898227. Host transcript counts are tabulated in (**Table S9**).

